# *Zbtb16* regulates social cognitive behaviors and neocortical development

**DOI:** 10.1101/2020.08.09.233270

**Authors:** Noriyoshi Usui, Stefano Berto, Ami Konishi, Makoto Kondo, Genevieve Konopka, Hideo Matsuzaki, Shoichi Shimada

**Affiliations:** Department of Neuroscience and Cell Biology, Graduate School of Medicine, Osaka University, Osaka 565-0871, Japan; Center for Medical Research and Education, Graduate School of Medicine, Osaka University, Osaka 565-0871, Japan; Global Center for Medical Engineering and Informatics, Osaka University, Osaka, 565-0871, Japan; Addiction Research Unit, Osaka Psychiatric Research Center, Osaka Psychiatric Medical Center, Osaka 541-8567, Japan; Department of Neuroscience, University of Texas Southwestern Medical Center, Dallas, TX 75390, USA; Department of Child Development, United Graduate School of Child Development, Osaka University, Osaka 565-0871, Japan; Division of Development of Mental Functions, Research Center for Child Mental Development, University of Fukui, Fukui 910-1193, Japan; Life Science Innovation Center, University of Fukui, Fukui, 910-1193, Japan

## Abstract

Recent genetic studies have underscored the pleiotropic effects of single genes to multiple cognitive disorders. Mutations of *ZBTB16* are associated with autism spectrum disorder (ASD) and schizophrenia (SCZ), but how the function of ZBTB16 is related to ASD or SCZ remains unknown. Here we show the deletion of *Zbtb16* in mice leads to both ASD- and SCZ-like behaviors such as social impairment, repetitive behaviors, risk-taking behaviors, and cognitive impairment. To elucidate the mechanism underlying the behavioral phenotypes, we carried out histological studies and observed impairments in thinning of neocortical layer 6 (L6) and a reduction of TBR1+ neurons in the prefrontal cortex (PFC) of *Zbtb16* KO mice. Furthermore, we found increased dendritic spines and microglia as well as developmental defects in oligodendrocytes and neocortical myelination in the PFC of *Zbtb16* KO mice. Using a genomics approach, we identified the *Zbtb16*-transcriptome that includes genes involved in both ASD and SCZ pathophysiology and neocortical maturation such as neurogenesis and myelination. Co-expression networks further identified *Zbtb16*-correlated modules that are unique to ASD or SCZ respectively. Our study provides insight into the differential role of *ZBTB16* in ASD and SCZ.

## Introduction

ASD is a heterogeneous neurodevelopmental disorder that causes pervasive abnormalities in social communication as well as repetitive behaviors and restricted interests. Worldwide, the prevalence of children diagnosed with ASD has increased significantly over recent decades^1^. The etiology of ASD is thought to involve complex, multigenic interactions, and possible environmental contributions^2^. SCZ is also a heterogeneous neuropsychiatric disorder characterized by positive symptoms (hallucinations and delusions), negative symptoms (flat affect, avolition, and social impairment), and cognitive impairment^3^. SCZ is a lifelong disorder that affects 1% of the world’s population and develops adolescence and young adulthood^4^. The biological mechanisms underlying ASD and SCZ are not fully understood. However, it is well-known that there are common overlapping mechanisms such as genetics, ethology, and brain dysfunction underlying the pathology of ASD and SCZ^5, 6^.

In this study, we aimed to uncover how ZBTB16 might be biologically relevant to both ASD and SCZ. The *ZBTB16 (PLZF*) gene is located in 11q23.2. A missense variant (c.1319G>A; p.Arg440Gln) of *ZBTB16* has been recently reported in brothers with ASD^7^. In contrast, a nonsense mutation in *ZBTB16* (c.1741A>T; p.Lys581*) has been also reported in SCZ patients^8, 9^. Moreover, there are additional reports on the association between *ZBTB16* and SCZ^10–12^. Duplication of 11q13.2-q25 where *ZBTB16* is located has been identified in children who had developmental delay with or without congenital malformations^13^. Other duplications of 11q22.1-q25 and 11q23.2-q23.3 have been also identified in individuals with ASD, and intellectual and developmental disabilities^14^.

*ZBTB16* encodes a transcription factor, which contains a BTB/POZ protein-protein interaction domain in its N-terminal and a C2H2-type zinc finger DNA binding domain in its C-terminal, playing key roles in many biological processes such as stem cell maintenance and proliferation, cell differentiation, spermatogenesis, musculoskeletal development, hematopoiesis, apoptosis, chromatin remodeling, metabolism and immunity^15, 16^. A variant translocation in the *ZBTB16* was first identified in a patient with acute promyelocytic leukemia^17^. Deletions of human 11q23 are associated with intellectual disability, microcephaly, epilepsy, craniofacial dysmorphism, and short stature^18–21^. Among the genes on 11q23, a single nucleotide variant (c.1849A>G) in the C2H2-type zinc finger domain of *ZBTB16* has been identified as a causative mutation for skeletal defects, genital hypoplasia, and mental retardation (SGYMR)^18, 22^. The SGYMR individual with this mutation in *ZBTB16* showed intellectual disability, microcephaly, craniofacial dysmorphism, short stature, skeletal anomalies such as thumb deficits and hypoplasia of the ulnae, retarded bone age, and hypoplastic genitalia^18, 22^. Studies of spontaneous *luxoid* (*lu*) mutant of *Zbtb16* in mice have shown that *Zbtb16* is essential for skeletal development and germ cell self-renewal^23–25^.

Involvement of *ZBTB16* in brain development has been reported in several studies. *Zbtb16* expression begins at embryonic day 7.5 in the neuroepithelium of the mouse embryonic brain, and is eventually expressed in the entire neurectoderm at later stages^26^. *ZBTB16* is expressed in human embryonic stem cell (ES)-derived neural stem cells (NSC) and primary neural plate tissue, playing a role in maintenance, proliferation^27^, and neuronal differentiation^28^. A recent study has reported reduced cortical surface area and a number of deep layer neurons in the neocortex of *Zbtb16^lu/lu^* mutant mice at neonatal stages^29^. In addition, *Zbtb16^lu/lu^* mutant mice showed an impairment of recognition memory in the novel object recognition test^29^. These studies indicate the involvement of *Zbtb16* in neocortical development. However, these previous findings were primarily limited to the embryonic and neonatal periods, whereas the role of *Zbtb16* in the adult brain, particularly association with multiple diseases such as ASD and SCZ are unknown.

Again, in this study, we aimed to uncover how *Zbtb16* regulates the pathological mechanisms underlying ASD and SCZ. To address this question, we utilized a *Zbtb16^lu^* homozygous mutant (*Zbtb16* KO) mouse, and characterized the behavioral and neocortical phenotypes. *Zbtb16* KO mice displayed an impairment in social behaviors and increased repetitive behaviors. *Zbtb16* KO mice also showed increased risk-taking behaviors and cognitive impairment. We analyzed the PFC of *Zbtb16* KO mice, and found that impairments in neocortical thickness and a reduction of TBR1+ neuronal numbers. We also found increases in the numbers of dendritic spines and microglia. Moreover, we found reductions in oligodendrocyte development, resulting in impaired neocortical myelination. Finally, we characterized the *Zbtb16* transcriptome in the PFC by conducting RNA-sequencing (RNA-seq), and identified that *Zbtb16* regulates genes known to be involved in ASD, SCZ, and neocortical maturation including myelination. We further investigated co-expression networks and identified the disorder-specific independent genes for ASD or SCZ respectively. These results demonstrate that *Zbtb16* plays an essential role in both ASD- and SCZ-like behaviors via neocortical development, particularly deep layer formation, spinogenesis, and myelination. Taken together, our study demonstrate that *Zbtb16* is involved in shared neurodevelopmental features that are at risk in both ASD and SCZ.

## Materials and Methods

### Mice

B6.C3-*Zbtb16^lu^*/J mice were purchased from The Jackson Laboratory (#000100)^23, 24^. Genotyping was performed using the following primers; for *Zbtb16*: 23559, F-5’-CCACCTCTTTCGGTCTCTCA-3’; 23560, R-5’-CCCCTCTTTGCTCCTCTCTT-3’ to detect a point mutation (C>T) by Sanger sequencing. Mice were housed in the barrier facilities of Osaka University under a 12 h light–dark cycle and given ad libitum access to water and food. All procedures were approved by the Animal Research Committee of Osaka University.

### Behavioral overview

All mice used for behavioral testing were 7-8 weeks male littermate progeny of heterozygous *Zbtb16* mutant crossings. An experimenter blind to genotypes performed all behavioral tests. The behavioral cohort consisted of the following age-matched littermate triplets and pairs: wild-type (WT)=18, KO=15 for stereotyped behavior, locomotion, open field, elevated plus maze, three-chamber social interaction, and marble-burying tests; WT=14, KO=8 for novel object recognition test. All behavioral tests were performed between 1000 to 1600 hours.

### Stereotyped behavior test

Mice were placed in a novel home cage where they were habituated for 10 min followed by a 10 min recording period. Time spent and number of grooming events were manually quantified from videotape.

### Locomotion test

Mice were placed in a novel chamber (W700 × D700 × H400 mm, #OF-36(M)SQ, Muromachi Kikai Co., Ltd., Tokyo, Japan) and allowed to freely explore for 10 min. Horizontal locomotor activity was measured by ANY-maze behavior tracking software (Stoelting Co., Wood Dale, IL). Habituation to a novel camber was performed the day before the test.

### Open field test

Mice were placed in one of the corners of a novel chamber (W700 × D700 × H400 mm, #OF-36(M)SQ, Muromachi Kikai Co., Ltd., Tokyo, Japan) and allowed to freely explore for 10 min. Time spent in the center of the arena (140 × 140 mm) and in all corners of the arena (140 × 140 mm × 4 corners), and locomotor activity were measured and tracked by ANY-maze behavior tracking software.

### Elevated plus maze test

Mice were placed in the center of the maze (open arms W54 × D297 mm; closed arms W60 × D300 × H150 mm; Height from floor 400 mm, # EPM-04M, Muromachi Kikai Co., Ltd., Tokyo, Japan) and allowed to freely explore the maze for 5 min. Time and distance in each arm were measured and tracked by ANY-maze behavior tracking software.

### Three-chamber social interaction test

Social interaction test consisted of three 5 min trials in the three-chamber apparatus (W600 × D400 × H220 mm, SC-03M, Muromachi Kikai Co., Ltd., Tokyo, Japan). During the first trial, the mouse was allowed free exploration of the three-chamber apparatus. Each end chamber contained an empty wire cage (φ90 × H185 mm) with the middle chamber being empty. In the second 5 min trial to examine social novelty, one of the end chambers contained a novel stranger mouse in a wire cage while the opposite end chamber contained an empty wire cage. In the third 5 min trial to examine social cognition, one of the end chambers kept a mouse used in second trial in same wire cage as a familiar mouse, while the opposite end chamber contained the other novel stranger mouse in a wire cage. The test mouse was also given a choice between an inanimate cage and a novel stranger mouse in the second trial, and a familiar mouse and a novel stranger mouse in the third trial. Interaction with the targets around a wire cage was tracked and measured by ANY-maze behavior tracking software.

### Marble-burying test

Mice were placed in the corner of a novel home cage evenly placed eighteen marbles and allowed to freely explore for 20 min. After 20 min, the number of marbles buried was recorded. A marble was defined as buried when less than one-third of the marble was visible.

### Novel object recognition test

Mice were habituated the day before the test to a chamber (W700 × D700 × H400 mm, #OF-36(M)SQ, Muromachi Kikai Co., Ltd., Tokyo, Japan). On the second day, two same objects were placed on the two opposite corners of a chamber from approximately 50 mm from the closest wall. Then, mice were placed in the corner of a chamber and allowed to freely explore for 10 min. On the third day, one of the objects was replaced to a different shaped object as a novel object. Mice were placed in the corner of a chamber and allowed to freely explore for 10 min. Interaction with novel and familiar objects was tracked and measured by ANY-maze behavior tracking software. The difference score was calculated by subtracting the time exploring the familiar object from the time exploring the novel object. The discrimination ratio was calculated by dividing the time exploring the familiar object by the total time exploring both novel and familiar objects. A positive difference score or discrimination ratio >0.5 indicates that a mouse recognizes the novel object.

### Immunohistochemistry

Mouse brains at 7-8 weeks were fixed with 4% PFA in PBS overnight at 4°C, cryoprotected in 30% sucrose in PBS, then embedded in Tissue-Tek O.C.T. Compound (#4583, Sakura Finetek Japan Co., Ltd., Osaka, Japan) for cryosectioning. Cryosections (20 μm thick) were placed in PBS. Antigen retrieval pretreatment was performed by incubating sections in citrate buffer (10 mM citrate, 0.05% Tween-20, pH 6) at 95°C for 10 minutes. Sections were stained with the following primary antibodies: mouse monoclonal anti-NeuN (1:200, #MAB377, Millipore, Billerica, MA), rat monoclonal anti-CTIP2 (1:500, #ab18465, Abcam, Cambridge, UK), rabbit polyclonal anti-TBR1 (1:250, #ab31940, Abcam, Cambridge, UK), mouse monoclonal anti-PDGFRα (CD140a) (1:200, #14-1401-82, Thermo Fisher Scientific, Waltham, MA), mouse monoclonal anti-APC (1:250, #OP80, Merck, Darmstadt, Deutschland), rabbit polyclonal anti-MBP (1:200, #ab40390, Abcam, Cambridge, UK), rabbit polyclonal anti-IBA1 (1:1000, #019-19741, FUJIFILM Wako pure chemical corporation, Osaka, Japan). For fluorescence immunostaining, species-specific antibodies conjugated to Alexa Fluor 488 and/or Alexa Fluor 597 (1:2,000; Invitrogen, Carlsbad, CA) were applied, and cover glasses were mounted with Fluoromount/Plus (#K048, Diagnostic BioSystems, Pleasanton, CA) or ProLong Diamond Antifade Mountant with DAPI (#P-36931 or #P36971, Thermo Fisher Scientific, Waltham, MA) for nuclear staining. DAPI (#11034-56, Nacalai Tesque, Kyoto, Japan) was also used to stain the nucleus. Images were collected using Olympus microscope and digital camera system (BX53 and DP73, Olympus, Tokyo, Japan) and All-in-One fluorescence microscope (BZ-X700, KEYENCE Corporation, Osaka, Japan). Cell numbers and cortical thickness (at bregma −1.70 mm) were quantified manually or using KEYENCE analysis software with Hybrid cell count application (KEYENCE Corporation, Osaka, Japan). Myelinated cortical area (at bregma −1.70 mm) was quantified as described previously^30^.

### Golgi staining

Whole brains collected at 7-8 weeks were subjected to Golgi staining using the superGolgi Kit (#003010, Bioenno Tech, LLC, Santa Ana, CA) according to the manufacturer’s instruction. Coronal sections (100 μm thick) were cut using a vibrating blade microtome (VT1000S, Leica Biosystems, Wetzlar, Germany), and mounted on the slides. Images were collected using KEYENCE analysis software with quick full focus (KEYENCE Corporation, Osaka, Japan).

### RNA-sequencing (RNA-seq)

RNA-seq was performed as a service by Macrogen Japan Corp. (Kyoto, Japan). Briefly, total RNA was extracted from prefrontal cortex (PFC) of male mice at 7 weeks with the AllPrep DNA/RNA Mini Kit (#80204, Qiagen, Hilden, Germany) according to the manufacturer’s instruction. RNA integrity number (RIN) of total RNA was quantified by Agilent 2100 Bioanalyzer using Agilent RNA 6000 Pico Kit (#5067-1513, Agilent, Santa Clara, CA). Total RNA with RIN values of ≥8.1 were used for RNA-seq library preparation. mRNA was purified from 500 ng total RNA, and subjected to cDNA library making (fragmentation, first and second strand syntheses, adenylation, ligation and amplification) by TruSeq Stranded mRNA Library Prep (#20020594, Illumina, San Diego, CA) according to the manufacturer’s instruction. cDNA library quality was quantified by 2100 Bioanalyzer using Agilent High Sensitivity DNA Kit (#5067-4626, Agilent, Santa Clara, CA). Library was sequenced as 101 bp paired-end on Illumina NovaSeq6000.

### RNA-seq alignment and quality control

Reads were aligned to the mouse mm10 reference genome using STAR (v2.7.1a)^31^. For each sample, a BAM file including mapped and unmapped reads that spanned splice junctions was produced. Secondary alignment and multi-mapped reads were further removed using in-house scripts. Only uniquely mapped reads were retained for further analyses. Quality control metrics were assessed by Picard tool (http://broadinstitute.github.io/picard/). Gencode annotation for mm10 (version M21) was used as reference alignment annotation and downstream quantification. Gene level expression was calculated using HTseq (v0.9.1)^32^ using intersection-strict mode by exon. Counts were calculated based on protein-coding genes from the annotation file.

### Differential expression

Counts were normalized using counts per million reads (CPM). Genes with no reads in either *Zbtb16* KO or WT samples were removed. Surrogates variables were calculated using *sva* package in R^33^. Differential expression analysis was performed in R using linear modeling as following: lm(gene expression ~ Treatment + nSVs). We estimated log2 fold changes and P-values. P-values were adjusted for multiple comparisons using a Benjamini-Hochberg correction (FDR). Differentially expressed genes where consider for FDR<0.05. Mouse Gene ID were translated into Human Gene ID using *biomaRt* package in R^34^.

### Gene ontology analyses

The functional annotation of differentially expressed and co-expressed genes was performed using GOstats^35^. A Benjamini-Hochberg FDR (FDR<0.05) was applied as a multiple comparison adjustment.

### Network analyses

We carried out weighted gene co-expression network analysis (WGCNA)^36^. Prior the co-expression analysis, normalized RNA-seq data was residualized and balanced for the nSVs detected using a linear model. A soft-threshold power was automatically calculated to achieve approximate scale-free topology (R2>0.85). Networks were constructed with blockwiseConsensusModules function with biweight midcorrelation (bicor). We used corType = bicor, networkType = signed, TOMtype = signed, TOMDenom = mean, maxBlockSize = 16000, mergingThresh = 0.15, minCoreKME = 0.5, minKMEtoStay = 0.6, reassignThreshold = 1e-10, deepSplit = 4, detectCutHeight = 0.999, minModuleSize = 50. The modules were then determined using the dynamic tree-cutting algorithm. Deep split of 4 was used to more aggressively split the data and create more specific modules. Spearman’s rank correlation was used to compute module eigengene – treatment association.

### GWAS data and enrichment

We used genome-wide gene-based association analysis implementing MAGMA v1.07^37^. We used the 19346 protein-coding genes from human gencode v19 as background for the gene-based association analysis. SNPs were selected within exonic, intronic, and UTR regions as well as SNPs within 5kb up/down-stream the protein-coding gene. SNP association revealed 18988 protein-coding genes with at least one SNP. Gene based association test were performed using linkage disequilibrium between SNPs. Benjamini-Hochberg correction was applied and significant enrichment is reported for FDR<0.05. Summary statistics for GWAS studies on neuropsychiatric disorders and non-brain disorders were downloaded from Psychiatric Genomics Consortium and GIANT Consortium^38–48^. GWAS acronyms were used for the figures (*ADHD = attention deficit hyperactivity disorder, AZ = Alzheimer’s disease, ASD = autism spectrum disorder, BD = bipolar disorder, Epilepsy = epilepsy, MDD = major depressive disorder, SCZ = schizophrenia, CognFunc = cognitive functions, EduAtt = educational attainment, Intelligence = Intelligence, BMI = body mass index, CAD = coronary artery disease, OSTEO = osteoporosis*).

### Gene set enrichment

Gene set enrichment was performed in R using Fisher’s exact test with the following parameters: alternative = “greater”, confidence level = 0.95. We reported Odds Ratio (OR) and Benjamini-Hochberg adjusted P-values (FDR).

### Statistical analysis

All behavioral and histological data are represented as means of biological independent experiments with ±standard error of the mean (sem). Statistical analysis (unpaired *t*-test) was performed using Prism 7. Asterisks indicate P-values (****P<0.0001, ***P<0.001, **P<0.01, *P<0.05).

## Results

### *Zbtb16* KO mice display ASD-like behaviors

*Zbtb16* KO mice showed skeletal dysplasia and smaller body size compared with WT mice (Fig. 1a), due to a single nucleotide (*lu*) mutation in *Zbtb16*, resulting in a nonsense mutation (p.Arg234*) (Fig. 1b, Supplementary Figure 1). We first investigated locomotion activity in the mice. *Zbtb16* KO mice showed weight loss (P<0.0001) (Fig. 1c), but no difference in normal locomotion activity (P=0.8664) (Fig. 1d).

**Figure 1.**
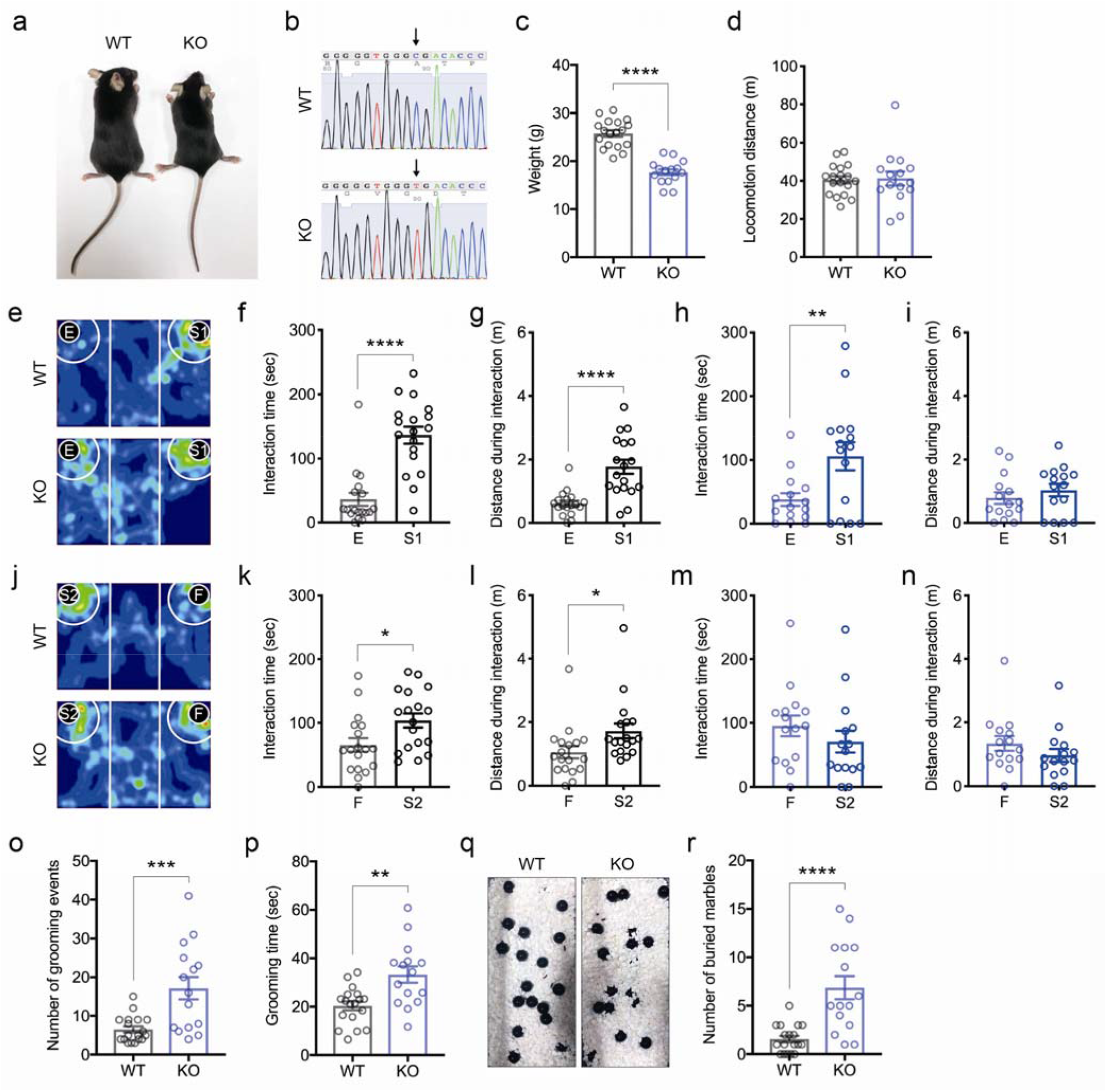
*Zbtb16* KO mice display social impairment and repetitive behaviors. **a** *Zbtb16* KO mice display small bodies and hind limb dysplasia. *WT: wild-type, KO: Zbtb16 knockout*. **b** A single nucleotide (*luxoid*) mutation in *Zbtb16* results in a nonsense mutation (p.Arg234*). **c** Weight loss in *Zbtb16* KO mice. **d** No serious impairments in locomotion activity in *Zbtb16* KO mice. **e** Representative heatmaps of social novelty behavior in the 3-chamber social interaction test. *E: empty, S1: stranger mouse 1*. **f,g** Quantification of interaction time (**f**) and distance during interaction (**g**) in social novelty section in WT mice. **h,i** Quantification of interaction time (**h**) and distance during interaction (**i**) in social novelty section in *Zbtb16* KO mice. **j** Representative heatmaps of social cognitive behavior in the 3-chamber social interaction test. *S2: stranger mouse 2, F: familiar mouse*. **k,l** Quantification of interaction time (**k**) and distance travelled during interaction (**l**) in social cognition section in WT mice. **m,n** Quantification of interaction time (**m**) and distance travelled during interaction (**n**) in social cognition section in *Zbtb16* KO mice. WT mice spent more time and distance interacting with a stranger mouse than a familiar mouse, compared to *Zbtb16* KO mice. **o** Quantification of grooming events. **p** Quantification of the number of grooming times. **q** Representative images after a marble-burying test. **r** Quantification of the number of buried marbles. *Zbtb16* KO mice showed an increase in repetitive behaviors like burying marbles. Data are represented as means (±SEM). ****P<0.0001, ***P<0.001, **P<0.01, *P<0.05, unpaired t-test. n =15-18/condition.

Since there was no serious problem in the locomotion activity of *Zbtb16* KO mice that could confound conducting behavioral analyses (Fig. 1d), we investigated whether *Zbtb16* KO mice exhibit behaviors relevant to ASD. We first examined social behaviors using a three-chamber social interaction test. In the social novelty trial, WT mice preferred a novel mouse than an inanimate empty cage, and spent more time (P<0.0001) and distance (P<0.0001) interacting with a social target (Fig. 1e-g). On the other hand, *Zbtb16* KO mice also preferred a novel mouse over an inanimate empty cage; however, *Zbtb16* KO mice spent increased time (P=0.0090), but not distance (P=0.3675) interacting with a social target (Fig. 1e, i, h). In the social cognition trial, WT mice spent more time (P=0.0178) and distance (P=0.0322) interacting with a novel mouse than a familiar mouse (Fig. 1j-l). In contrast, *Zbtb16* KO mice spent approximately similar time (P=0.3059) and distance (P=0.2389) for interacting with both a novel mouse and a familiar mouse (Fig. 1j, m, n). These results indicate that *Zbtb16* KO mice show decreased social novelty and impaired social cognition.

We next examined repetitive behaviors, and found that the number of grooming events (P=0.0006) and grooming time (P=0.0014) were significantly increased in *Zbtb16* KO mice (Fig. 1o, p). In addition, we found that *Zbtb16* KO buried more marbles compared with WT mice (P<0.0001) (Fig. 1q, r). These results indicate that *Zbtb16* KO mice show increased repetitive behaviors.

Together, these results demonstrate that *Zbtb16* KO mice display ASD-relevant behaviors.

### *Zbtb16* KO mice show SCZ-like behaviors

We next investigated whether *Zbtb16* KO mice show behaviors relevant to SCZ. *Zbtb16* KO mice displayed social impairment, which is similar to one of the negative symptoms in SCZ. It has been reported that impulsive risk-taking behaviors are common in patients with SCZ^49, 50^. Thus, we analyzed risk-taking (antianxiety) behaviors in *Zbtb16* KO mice using an open field test^51^. We found that *Zbtb16* KO mice spent more time (P=0.0217) and distance (P=0.0468), but not the number of entries (P=0.0784) in the center of the arena compared to WT mice (Fig. 2a-d). On the other hand, there were no differences in the number of entry (P=0.6939), time (P=0.2738), distance (P=0.6349) in the corners of the arena (Fig. 2a, e-g). These results indicate that *Zbtb16* KO mice explore the field more than WT mice, suggesting risk-taking behavior is increased in *Zbtb16* KO mice. To clarify the risk-taking behavior of *Zbrb16* KO mice, we performed elevated plus maze^51^. Similarly, we found significantly increased time (P=0.0004) and distance (P=0.0048), but not the number of entry (P=0.3391) in open arms of maze in *Zbtb16* KO mice compared with WT mice (Fig. 2h-k). In addition, we found significantly decreased time (P=0.0035), but not the number of entry (P=0.4090) and distance (P=0.4966) in closed arms of maze in *Zbtb16* KO mice. These results demonstrate that *Zbtb16* KO mice show risk-taking behaviors.

**Figure 2.**
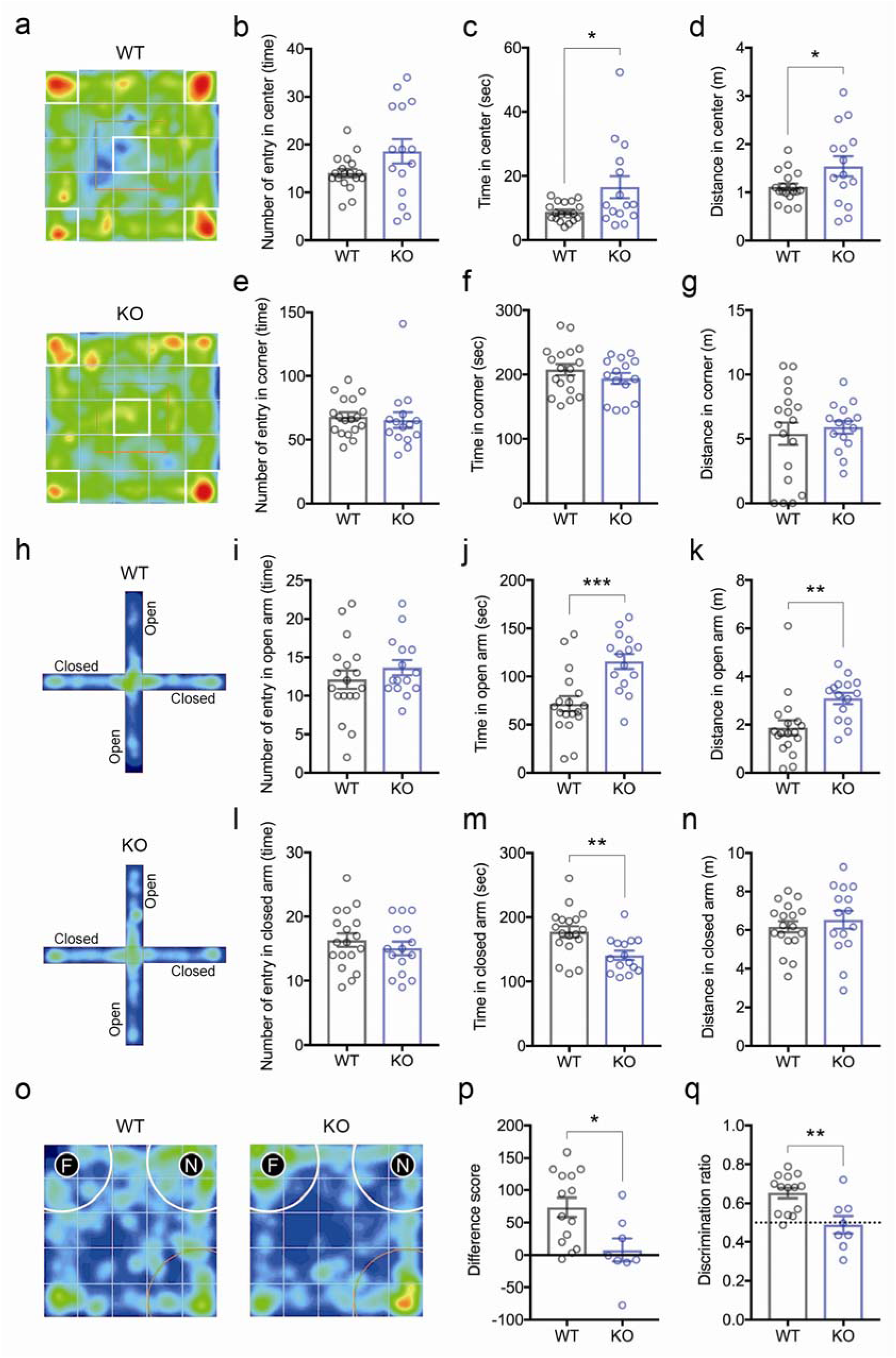
*Zbtb16* KO mice show risk-taking behaviors due to cognitive impairment. **a** Representative heatmaps of anxiety-like behavior in the open field test. **b-g** Quantification of the number of entries in center (**b**), time in center (**c**), distance in center (**d**), the number of entries in corner (**e**), time in corner (**f**), and distance in corner (**g**) during the open field test. *Zbtb16* KO mice spent more time in center than WT mice. **h** Representative heatmaps of anxiety-like behavior in the elevated plus maze test. *N: Novel mouse, F: familiar mouse*. **i-n** Quantification of the number of entries in open arm (**i**), time in open arm (**j**), distance in open arm (**k**), the number of entries in closed arm (**l**), time in closed arm (**m**), and distance in closed arm (**n**) during the elevated plus maze test. *Zbtb16* KO mice also spent more time in the open arm than closed arm compared with WT mice, indicating that impulsive risk-taking behaviors were increased in *Zbtb16* KO mice. **o** Representative heatmaps of learning and memory behaviors in the novel object recognition test. **p-q** Quantification of difference score (**p**) and discrimination ratio (**q**) in the novel object recognition test. Cognitive function was significantly impaired in *Zbtb16* KO mice. Positive difference score or discrimination ratio >0.5 means a mouse recognizes the novel object. Data are represented as means (±SEM). ***P<0.001, **P<0.01, *P 0.05, unpaired t-test. n=15-18/condition for open field and elevated plus maze tests, n=8-14/condition for novel object recognition test.

Since cognitive impairment in the novel object recognition test in *Zbtb16* KO mice (without age information) has been reported^29^, we also investigated the cognitive function of *Zbtb16* KO mice in the same test. As we expected, we observed significant reductions in difference score (P=0.0122) and discriminant ratio (P=0.0032) of the novel object recognition test in *Zbtb16* KO mice (Fig. 2o-q). These results indicate that cognitive function, in particular learning and memory are impaired in *Zbtb16* KO mice.

Together, these results demonstrate that *Zbtb16* KO mice also display behaviors relevant to SCZ.

### Impairments of neocortical deep layer formation in *Zbtb16* KO mice

To place the behavioral abnormalities in a biological context, we analyzed the development of the neocortex of *Zbtb16* KO mice. *Zbtb16* is mainly expressed in the neocortex, striatum, amygdala, hippocampus, midbrain, and cerebellum (Supplementary Figure 2). Since reduced neocortical area and abnormal deep layer formation in neonatal *Zbtb16* KO mice has been reported^29^, we examined neocortical thickness and lamination in the PFC of *Zbtb16* KO mice at 7-8 weeks. The reduced brain size of *Zbtb16* KO mice was observed as previously reported (Fig. 3a). We quantified neocortical thickness with DAPI staining, and found that significant reduction of neocortical thickness in the PFC of *Zbtb16* KO mice (P=0.0087) (Fig. 3b, c). Moreover, we identified that L6 (P=0.0486) was specifically thinner in *Zbtb16* KO mice compared to WT mice, but not L5 (P=0.3897) as measured by immunostaining using CTIP2 and TBR1, deep layer markers for L5 and L6 respectively (Fig. 3d, e). We also found a significant reduction of TBR1+ cells (P<0.0001) in the PFC of *Zbtb16* KO mice, but not CTIP2+ cells (P=0.7202) (Fig. 3f, g). These results demonstrate that deep layer formation, in particular L6 formation was impaired in *Zbtb16* KO mice.

**Figure 3.**
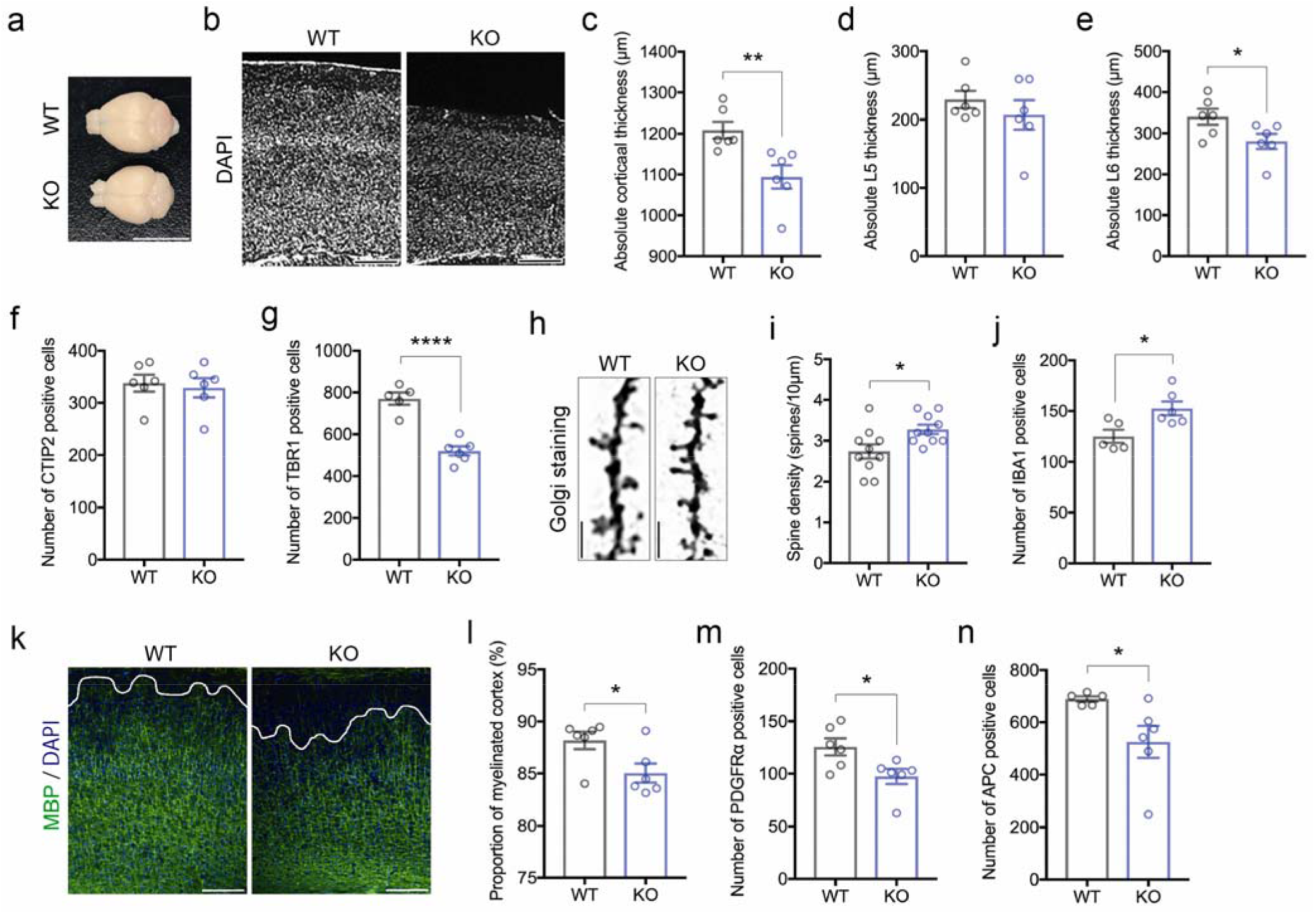
Impairments of neocortical development in *Zbtb16* KO mice. **a** *Zbtb16* KO mice show smaller brains than WT mice. **b** Representative images showing thinner neocortical thickness in *Zbtb16* KO mice. **c** Quantifications of neocortical thickness by DAPI staining. A significant reduction of neocortical thickness was observed in the PFC of *Ztbt16* KO mice. **d-e** Quantification of deep layer thickness by co-immunostaining of CTIP2, a marker for layer 5 (L5) (**d**) and TBR1, a marker for L6 (**e**). Neocortical L6 was specifically thinner in *Ztbt16* KO mice. **f-g** Quantification of CTIP2+ (**f**) and TBR1+ (**g**) cells in the mouse PFC. The number of TBR1+ L6 neurons was specifically decreased in the PFC of *Ztbt16* KO mice. **h** Representative fluorescent images of dendritic spines by Golgi staining. **i** Quantification of dendritic spines in the mouse PFC. **J** Quantification of IBA1+ microglia the mouse PFC. Significant increases of dendritic spines (**i**) and IBA1+ microglia (**j**) were observed in the PFC of *Ztbt16* KO mice. **k** Representative fluorescent images of myelination in the PFC by MBP immunostaining. **l** Quantification of proportion of myelinated neocortex. Immature myelin formation was observed in the PFC of *Ztbt16* KO mice. **m-n** Quantifications of oligodendrocyte in the PFC. Significant reductions of PDGFRα+ oligodendrocyte progenitors (**m**) and APC+ mature oligodendrocytes (**n**) were observed in the PFC of *Ztbt16* KO mice. Data are represented as means (±SEM). ****P<0.0001, **P<0.01, *P 0.05, unpaired t-test. n=5-6/condition. Scale bars: 1000 μm in **a**, 300 μm in **b**, **k**, 10 μm in **h**.

### Increased numbers of dendritic spines and microglia in *Zbtb16* KO mice

We further investigated dendritic spine and microglia, because it has been reported that the numbers of spines and microglia are linked to ASD and SCZ^52, 53^. Thus, we quantified the number of dendritic spines in the PFC of *Zbtb16* KO mice by Golgi staining. The number of dendritic spines was significantly increased in *Zbtb16* KO mice (P=0.0178) (Fig. 3h, i). Since microglia are responsible for spine pruning, we analyzed microglia in the PFC of *Zbtb16* KO mice by immunostaining. The number of IBA1+ microglia was significantly increased in *Zbtb16* KO mice (P=0.0175) (Fig. 3j). These results indicate that increased numbers of dendritic spines and microglia in *Zbtb16* KO mice also underlie functional abnormalities in the PFC.

### Immature myelination occurs due to oligodendrocyte loss in *Zbtb16* KO mice

Since myelination is associated with ASD and SCZ as well as cognitive functions^54^, we therefore examined myelination in the PFC of *Zbtb16* KO mice by immunostaining with MBP, a structural component of myelin. The ratio of myelinated area to the total area of the PFC was measured as previously reported^30^. The proportion of myelinated cortex was significantly decreased in *Zbtb16* KO mice (P=0.0304) (Fig. 3k, l). To clarify the cause of the myelination defect, we investigated oligodendrocyte development by immunostaining with oligodendrocyte differentiation markers, and found significant reductions of PDGFRα+ immature (P=0.0269) and APC+ mature (P=0.0409) oligodendrocytes in the PFC of *Zbtb16* KO mice (Fig. 5m, n). These results suggest that decreased myelination is due to abnormal oligodendrocyte development in *Zbtb16* KO mice.

Together, our results suggest that the histological abnormalities in neocortical cytoarchitectures such as L6 formation, dendritic spines, microglia, and myelination may underlie the behavioral deficits of *Zbtb16* KO mice.

### *Zbtb16* regulates neurodevelopmental genes and myelination-associated genes

To understand the molecular mechanisms underlying behavioral and histological phenotypes, we characterized the *Zbtb16* transcriptome by RNA-seq. Using PFC from 5 WT and 5 *Zbtb16* KO mice, we clearly separate the transcriptome profiles between WT and *Zbtb16* KO mice (Fig. 4a, Supplementary Figure 3). Differential expression analysis of the RNA-seq data uncovered 533 differentially expressed genes (DEGs) (FDR<0.05) in PFC of *Zbtb16* KO mice compared to WT mice (Fig. 4b, Supplementary Table 1). *Chrm2, Ddhd1, Dlat, Hspa4l, Igfbp3, Rnf169, Slc29a4, Vrk1, Zbtb16, and Fgf13* were identified as the top 10 DEGs in *Zbtb16* KO mice (Fig. 4b).

**Figure 4.**
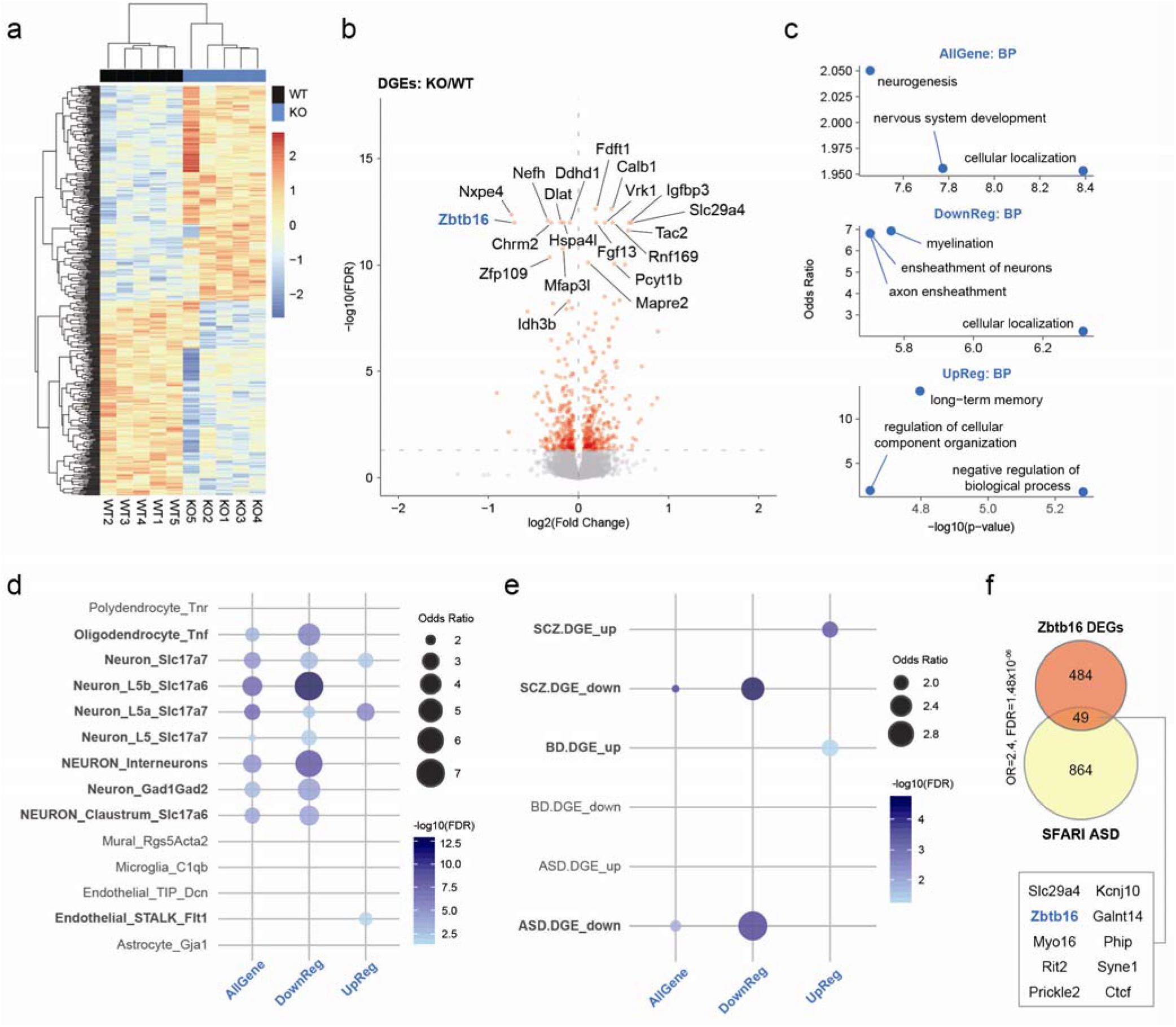
The *Zbtb16* transcriptome is enriched for ASD, SCZ and cortical development-related genes. **a** Heatmap of differential expressed genes (DEGs) in mouse PFC. **b** Volcano plot showing *Zbtb16* DEGs with the top 20 gene names indicated. FDR<0.05, Y-axis=-log10(FDR), X-axis=log2(Fold Change). **c** GO analyses of *Zbtb16* DEGs in biological process (BP). Scatterplots represent the top 3 functions in each module. Y-axis=Odds Ratio, X-axis=-log10(p-value). *AllGene: all DEGs, DownReg: downregulated DEGs, UpReg: upregulated DEGs*. **d** *Zbtb16* DEGs are enriched in neurons and oligodendrocyte. Dot plots of *Zbtb16* DEGs showing the cell-type specific enrichments. *Zbtb16* DEGs, in particular the downregulated DEGs are highly enriched in *Slc17a7+, Slc17a6+*, and *Gad1+ Gad2+* neurons, interneurons, and oligodendrocytes. In contrast, the upregulated DEGs are enriched in *Slc17a7+* neurons and endothelial cells. Dot plots represent Odds Ratio. Colors represent the −log10(FDR). **e** Dot plots of *Zbtb16* DEGs showing the enrichment for disorder-specific human transcriptome. Downregulated *Zbtb16* DEGs are enriched in ASD and SCZ downregulated genes. In contrast, upregulated *Zbtb16* DEGs are enriched in bipolar disorder and schizophrenia upregulated genes. *SCZ: schizophrenia, BD: bipolar disorder, ASD: autism spectrum disorder*. **f** Venn diagram of *Zbtb16* DEGs overlapping with SFARI ASD genes. The overlapping top 10 genes are highlighted.

We also performed gene ontology (GO) analysis to identify the functions of DEGs. DEGs are involved in cellular localization, nervous system development, and neurogenesis (Fig. 4c, Supplementary Figure 4). Interestingly, we found cellular localization, myelination, and axon ensheathment enriched among the downregulated DEGs (Fig. 4c, Supplementary Figure 4). The negative regulation of biological process, long-term memory, and regulation of cellular component organization were found in the upregulated DEGs (Fig. 4c). Together, these results suggest that *Ztbt16* regulated genes are involved in neurogenesis, myelination, and memory.

To further characterize DEGs, we conducted cell-type specific enrichment analysis using scRNA-seq data sets (see Methods) and *Zbtb16* transcriptome data. We found that the downregulated *Zbtb16* DEGs were highly enriched in *Slc17a7+, Slc17a6+*, and *Gad1+ Gad2+* neurons, and interneurons as well as oligodendrocytes (Fig. 4d, Supplementary Table 2). In contrast, the upregulated *Zbtb16* DEGs were enriched in *Slc17a7+* neurons and endothelial cells (Fig. 4d, Supplementary Table 2). These results suggest that *Zbtb16* plays a role in neurogenesis of the deep layers and oligodendrogenesis.

### Zbtb16-regulated genes are associated with ASD and SCZ

We next examined whether the DEGs are associated with human diseases by enrichment analyses using disorder-specific human transcriptomes^5^. We found that DEGs were enriched for ASD- and SCZ-specific downregulated DEGs (OR=1.9, FDR=3.1×10^−03^; OR=1.8 FDR=2.3×10^−05^, respectively), in particular downregulated DEGs were highly enriched in ASD- and SCZ-specific downregulated DEGs (OR=3.2, FDR=3.1×10^−06^; OR=2.6, FDR=1.2×10^−06^, respectively) (Fig. 4e). In contrast, upregulated DEGs were enriched in SCZ- and bipolar disorder (BD)-specific upregulated DEGs (OR=2.1, FDR=0.02; OR=2.0, FDR=6.9×10^−05^, respectively) (Fig. 4e). We also investigated how many DEGs overlapped with ASD genes from the SFARI database, and found that 49 DEGs (approximately 10% of the DEGs) (OR=2.4, FDR=1.48×10^−06^) overlapped with SFARI ASD genes (Fig. 4f). These analyses indicate that the Ztbt16-regulated transcriptome is related to both ASD and SCZ. These transcriptomic findings point to molecular mechanism that could underlie the behavioral and histological phenotypes of *Zbtb16* KO mice.

### Co-expressed gene networks of regulated by Zbtb16 are related to ASD and SCZ genes

To identify the individual molecular networks for unique roles of *Zbtb16*, we conducted co-expression networks analyses using the *Zbtb16* transcriptome. Weighted gene co-expression network analysis (WGCNA) identified 35 modules (Supplementary Table 1). Among these modules, the brown, lightyellow, and royalblue modules were associated with *Zbtb16* genotype (Fig. 5a-d). The brown module had *Zbtb16* as one of its hub genes (Fig. 5b). A hub gene is a key modulator in the co-expression network. The majority of hub genes in these three modules were also enriched in *Zbtb16* DEGs (Supplementary Figure 5).

**Figure 5.**
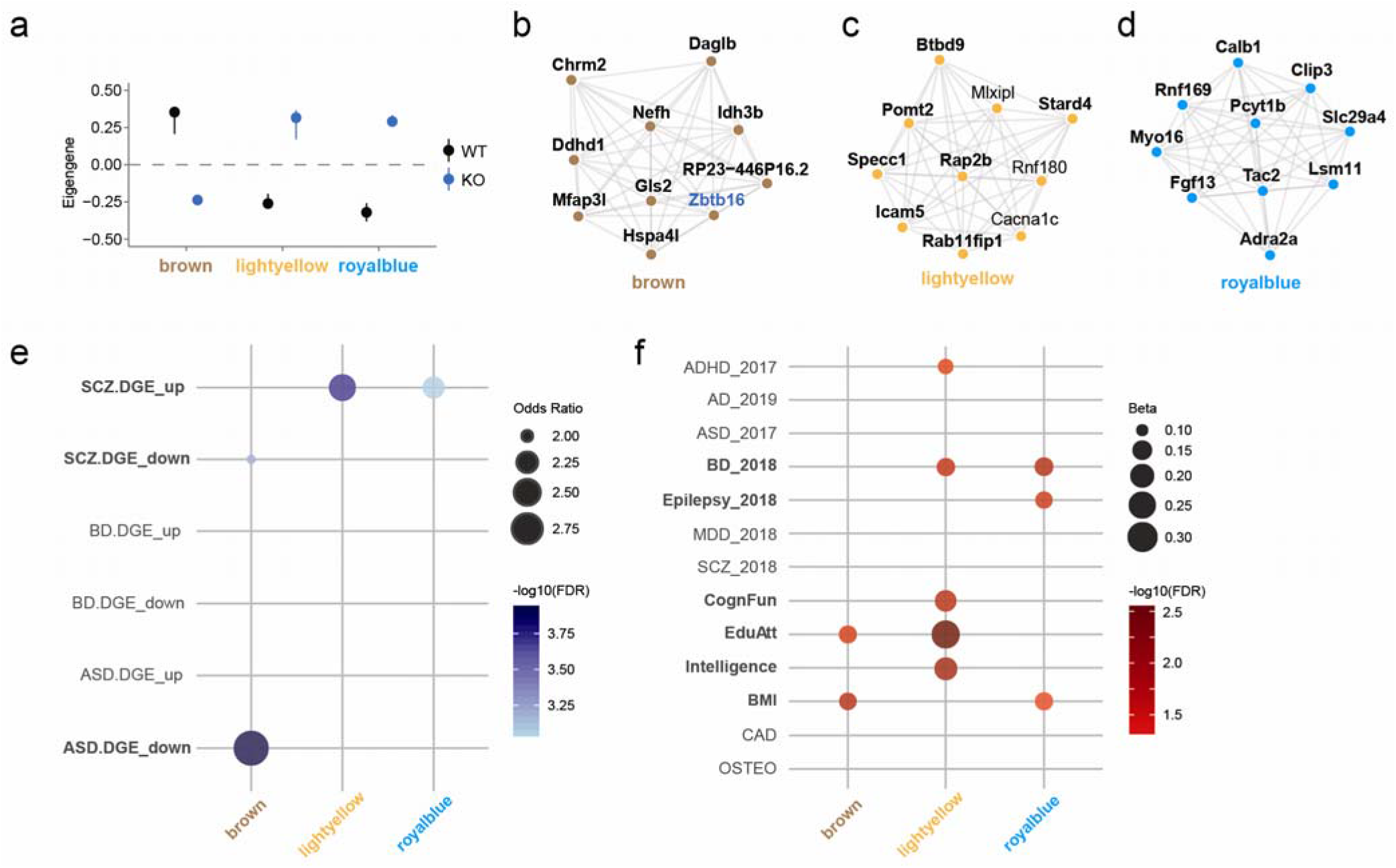
Co-expressed gene networks uncovered gene networks unique to ASD and SCZ. **a** Dot plots with standard errors (SEs) showing module eigengene demonstrate the association of the modules detected by parsimony for *Zbtb16*. Y-axis=rho, X-axis shows the modules highlighted. **b-d** *Zbtb16*-specific modules showing the top 10 hub genes ranked by weighted topological overlap values in the brown (**b**), lightyellow (**c**) and royalblue (**d**) modules. *Zbtb16* DEGs are shown in boldface. **e** *Zbtb16*-specific modules showing the enrichment for disorder-specific human transcriptome. The brown module is enriched for ASD and schizophrenia downregulated genes. The lightyellow and royalblue modules are enriched in schizophrenia upregulated genes. *SCZ: schizophrenia, BD: bipolar disorder, ASD: autism spectrum disorder*. **f** *Zbtb16*-specific modules are associated with BMI, cognitive traits, ADHD, BD and epilepsy. Dot plots represent Betas from MAGMA. Dot colors represent the −log10(FDR) from MAGMA. Y-axis shows the acronyms for the GWAS data utilized for this analysis (see Methods). X-axis shows the modules of this study. *ADHD: attention deficit hyperactivity disorder, AZ: Alzheimer’s disease, ASD: autism spectrum disorder, BD: bipolar disorder, Epilepsy: epilepsy, MDD: major depressive disorder, SCZ: schizophrenia*, *CognFunc: cognitive functions, EduAtt: educational attainment, Intelligence: Intelligence, BMI: body mass index, CAD: coronary artery disease, OSTEO: osteoporosis*.

We next examined whether the *Zbtb16*-specific modules show specific enrichment for ASD- or SCZ-related genes, and found that the brown module was enriched for ASD-specific downregulated DEGs (OR=2.9, FDR=6.5×10^−06^) (Fig. 5e). In contrast, the lightyellow and royalblue modules were enriched in SCZ-specific upregulated DEGs (OR=2.4, FDR=1.2×10^−05^; OR=2.2, FDR=1.9×10^−04^, respectively) (Fig. 5e). These results indicate that the brown module is highly enriched for ASD genes, and the other modules are enriched for SCZ genes. Finally, we examined whether the *Zbtb16*-specific modules were enriched for human genetic variants. The brown module was enriched in educational attainment (EduAtt) and body mass index (BMI) (FDR=0.03; FDR=0.02, respectively) (Fig. 5f, Supplementary Table 3). The lightyellow module was enriched in ADHD, BD, cognitive functions (CognFun), EduAtt, and intelligence (FDR=0.04; FDR=0.02; FDR=0.02; FDR=0.003; FDR=0.01, respectively) (Fig. 5f, Supplementary Table 3). The royalblue module was enriched in BD, epilepsy and BMI (FDR=0.01; FDR=0.02; FDR=0.05, respectively) (Fig. 5f, Supplementary Table 3). Our results demonstrate that these *Zbtb16*-specific modules are key gene networks for uncovering the molecular mechanisms of brain development at risk in both ASD and SCZ.

## Discussion

In this study, we demonstrate a role for *Zbtb16* in ASD- and SCZ-like behaviors such as social impairment, repetitive behaviors, risk-taking behaviors, and cognitive impairment. We also found histological impairments in cortical thickness and a reduction of TBR1+ neuronal numbers in the PFC of *Zbtb16* KO mice. Additionally, we found increases in the numbers of dendritic spines and microglia as well as defects in oligodendrocyte development and neocortical myelination in the PFC of *Zbtb16* KO mice. Transcriptomic analyses in the PFC of *Zbtb16* KO mice identified 533 DEGs involved in ASD, SCZ, and neocortical maturation such as neurogenesis and myelination. Co-expression networks analyses suggest that *Zbtb16*-specific modules may be distinct with respect to biological pathways underlying ASD and SCZ.

Associations of *ZBTB16* with ASD and SCZ have been reported. Here we uncovered behavioral signatures of *Zbtb16* KO mice (Fig. 1, 2). *Zbtb16* KO exhibited abnormal behaviors relevant to the core symptoms of ASD, social impairment and repetitive behaviors (Fig. 1), that are also consistent with the reported symptoms in the brothers with ASD and *ZBTB16* mutations^7^. On the other hand, *Zbtb16* KO mice showed social and cognitive impairment as well as risk-taking behaviors, which are all behaviors relevant to SCZ (Fig. 2). Risk-taking behavior is a key component not only in neuropsychiatric disorders such as SCZ^50, 55^, bipolar disorder^56^, and attention deficit hyperactivity disorder (ADHD)^57, 58^, but also in drug^59^ and alcohol^60^ abuse, and all of these involve dopamine signaling^61, 62^. The association between *Zbtb16* and neuropsychiatric disorders is further supported by the gene expression profiles in *Zbtb16* KO mice (Fig. 4, 5). For example, loss of the L6-specific gene *Foxp2* implicated in both ASD and SCZ in the cortex showed reduction of dopamine receptor D1 (Drd1), and these mice also displayed impaired behavioral flexibility^63^, suggesting that L6 is crucial for dopamine signal transduction and cognitive function. Consequentially, it is suggested that dopamine signaling might be altered in *Zbtb16* KO mice. In addition, the PFC plays the roles in short-term and long-term memory coordinating with hippocampus, and decision making^64^. Similarly, L6 neurons of visual cortex are also important for the processing the object-recognition memory^65^. Together, our results suggest that an increase in the risk-taking behaviors in *Zbtb16* KO mice is due to cognitive impairment. It will be interesting to explore the association of *Zbtb16* with dopamine signals and the roles of *Zbtb16* in the risk-taking behavior and cognitive function as future studies.

Previous studies have focused on the role of Zbtb16 in the maintenance and proliferation of NSC in embryonic stages^27, 28, 66^ and cortical surface area and thickness in neonatal stages^29^. Compared to those studies, we characterized the role of *Zbtb16* in the young adult brain, and found significantly decreased neocortical thickness of L6 and the number of TBR1+ cells (Fig. 3c, e, g) in *Zbtb16* KO mice^29^. Previous work has reported that a smaller cortex and loss of TBR1+ cells in *Zbtb16^lu/lu^* mutant mice are due to a decrease in proliferating mitotic cell numbers at early embryonic stages^29^. However, the mechanism that affects only deep neurons has not been clarified. Interestingly, it has been reported that the genes associated with ASD are enriched in the deep layers^67, 68^.

We also found increased numbers of dendritic spines and IBA1+ microglia in *Zbtb16* KO mice (Fig. 3h-j). The number of spines is well studied in neuropsychiatric disorders. It is frequently reported that spines increase in ASD and decrease in SCZ, but opposite phenotypes have also been reported^53, 69^. Changes in the number of spines are not only driven by genetic factors, but pruning by microglia is also an important factor. An increased number of microglia in postmortem brain^70, 71^ and activated intracerebral microglia^72, 73^ have been reported in patients with ASD. Microglial autophagy plays an essential role in dendritic spine pruning and social behaviors^74^. In summary, these findings suggest that an increase in the number of dendrite spines in PFC in *Zbtb16* KO mice could be a responsible factor for the observed impairment of social behavior.

Moreover, we found developmental defects of oligodendrocytes, resulting in impaired neocortical myelination in *Zbtb16* KO mice (Fig. 3k, l). This myelination-relevant phenotype was predicted by the RNA-seq results (Fig. 4c). To explain the impairment of neocortical myelination, we identified abnormal oligodendrocyte development via the reduced numbers of PDGFRα+ immature and APC+ mature oligodendrocytes in *Zbtb16* KO mice (Fig. 3m, n). *Zbtb16* is expressed in the OLIG2+ neural progenitor cells in early developmental stages, and regulates oligodendrocyte differentiation by suppressing neurogenesis^66^. Thus, these data suggest that the deletion of *Zbtb16* results in abnormal oligodendrogenesis, and eventually causes behavioral abnormalities such as ASD- and SCZ-like phenotypes. A recent study examined the role of the ASD-relevant gene TCF4 and found that *Tcf4* mutant mice show impairments of oligodendrocyte development and myelination, supporting the significance of oligodendrocytes implicating ASD etiology^75^. In addition, reductions in white matter or corpus callosum volumes in the brains of ASD^76, 77^ and SCZ^78, 79^ subjects using diffusion tensor imaging (DTI) have been reported. Collectively, our study demonstrates a novel role for *Zbtb16* in neocortical development such as abnormalities in deep layer formation, spinogenesis, and oligodendrogenesis, which are similar to both ASD and SCZ pathology.

Here, we describe the *Zbtb16* transcriptome in the adult mouse brain, which is also a resource for understanding the role of *Zbtb16* in brain development, behavior, and disease. In the RNA-seq results, we identified 533 DEGs in *Zbtb16* KO mice involved in essential biological functions such as neurogenesis, myelination, and memory (Fig. 3). Overlapping the *Zbtb16* DEGs with the list of the SFARI ASD genes (Fig. 4f), also provides ASD-relevant *Zbtb16*-mediated signaling pathways. Among 533 DEGs, downregulated targets of *Zbtb16* were significantly enriched for ASD- and SCZ-specific downregulated DEGs (Fig. 4d). For example, *CHRM2* plays a role in the communication between the cortex and hippocampus^80^ to modulate cognitive functions such as behavioral flexibility, working memory, and hippocampal plasticity^81^. A recent study has reported that variants in *DDHD1* are associated with hereditary spastic paraplegia and ASD^82^, and patients show clinical features such as cerebellar impairment, axonal neuropathy, distal sensory loss, and/or mitochondrial impairment^83, 84^. *HSPA4L* is significantly downregulated in lymphoblastoid cells from patients with SCZ^85^, however the function of HSPA4L in the brain is unknown. A recent in silico study has suggested that *HSPA4L* is upregulated in the corpus callosum of patients with multiple sclerosis, and is associated with myelination and the immune system^86^.

Lastly, we identified the ASD- (brown) and SCZ-specific (lightyellow and royalblue) *Zbtb16* modules using WGCNA (Fig. 5). In the brown module, the top hub genes (*Chrm2, Ddhd1, Zbtb16, Hspa4l, Mfap3l, Idh3b, Nefh, Gls2*, and *RP23-446P16.2*) were enriched for genes severely downregulated in ASD patients (Fig. 5b, e). Moreover, the brown module is enriched for genetic variants associated with EduAtt and BMI, and involved in mitochondrial functions (Supplementary Figure 6). In contrast, the lightyellow and royalblue modules showed specific enrichment for SCZ downregulated genes (Fig. 5c, e). At the genetic level, the lightyellow module is enriched for genetic variants associated with ADHD, BD, CognFun, EduAtt, and intelligence (Fig. 5c, f). The hub genes (*Rap2b, Btbd9, Pomt2, Specc1, Icam5, Rab11fip1, Stard4, Cacna1c, Rnf180*, and *Mlxipl*) of the lightyellow module are involved in behavior, memory, and synaptic membrane, further confirming the role of *Zbtb16* in these synaptic etiologies (Supplementary Figure 6). Furthermore, the royalblue module is enriched for genetic variants associated with epilepsy and BD (Fig. 5c, f). The hub genes (*Calb1, Clip3, Rnf169, Pcyt1b, Myo16, Slc29a4, Fgf13, Tac2, Lsm11*, and *Adra2a*) of the royalblue module are involved in dendritic spine, GTP binding, and purine ribonucleoside binding (Supplementary Figure 6). Consequently, detailed analyses of these *Zbtb16* target genes will lead to a deeper understanding that how *Zbtb16* regulates the pathological mechanism underlying ASD, SCZ and other neuropsychiatric disorders.

In summary, our study demonstrates a role for *Zbtb16* in social, repetitive, and risk-taking behaviors and cognitive function via neocortical development, particularly deep layer formation and myelination. To the best of our knowledge, this is the first study to directly show the association of *Zbtb16* with ASD- and SCZ-like behaviors as well as neocortical phenotypes and genomic associations. Our genomics results suggest that *ZBTB16* potentially regulates different gene networks that are at risks in ASD or SCZ. Further analysis of the mutations of *ZBTB16* (p.Arg440Gln in ASD and p.Lys581* in SCZ) would be the key to unlocking such molecular mechanisms. Presumably, alterations in protein conformation could play a role in the interaction of ZBTB16 with different binding factors, leading to differential downstream transcriptional targets. Further understanding of the differential roles of *Zbtb16* in the brain should give rise to novel insights and targets for understanding the molecular mechanisms underlying ASD and SCZ.

## Acknowledgements

We thank Yoko Sasaki, Ryoko Aramaki, Yuta Ono, Yuuto Ohara, Yuuki Takaba, Xie Min-Jue, and Tomoko Taniguchi for their support. This work was supported by the JSPS Grant-in-Aid for Scientific Research (C) (20K06872) to N.U.; JSPS Grant-in-Aid for Early-Career Scientists (18K14814) to N.U.; Takeda Science foundation to N.U.; SENSHIN Medical Research Foundation to N.U.; The Osaka Medical Research Foundation for Intractable Diseases to N.U.; Research Grant for Public Health Science to N.U.; Eli Lilly Japan Research Grant to N.U.; the Grant for Life Cycle Medicine from Faculty of Medical Sciences, University of Fukui to N.U.

## Author Contributions

N.U. designed the study, analyzed the data, and wrote the paper. N.U., A.K., and M.K. performed experiments and quantifications. S.B. and G.K. performed bioinformatic analyses. H.M. and S.S. supervised this study, and provided intellectual guidance. All authors discussed the results and commented on the manuscript.

