## Supplementary information for "*Zbtb16* regulates social cognitive behaviors and neocortical development"

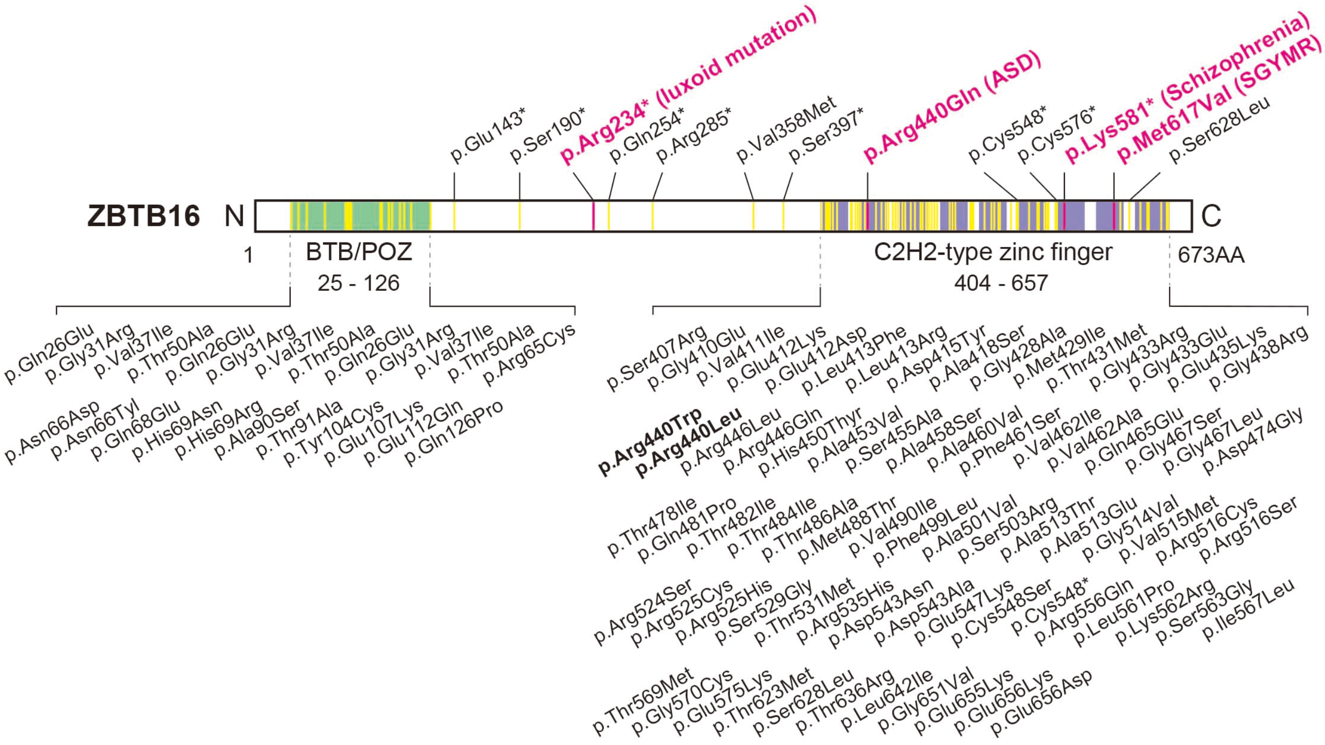


**Supplementary Figure 1.** ***Zbtb16*** **mutations and protein domains.**
Schematic of ZBTB16 protein showing the location of human mutations. The *ZBTB16* gene is located in 11q23.2. Duplications of 11q13.2-q25 have been identified in children who have developmental delay with or without congenital malformations[1]. Other duplications of 11q22.1-q25 and 11q23.2-q23.3 have also been identified in ASD, and intellectual and developmental disabilities[2]. A single nucleotide polymorphism (SNP) within the *ZBTB16* gene (c.1453+15115C>T; c.1454-7218C>T) showed association in the secondary analyses in a combined The Autism Genome Project (AGP) GWAS samples[3]. A spontaneous *luxoid* (*lu*) mutation (p.Arg234*) was identified in the *Zbtb16*$^{lu}$ mutant mouse[4, 5], which is highly conserved across humans to zebrafish. A SNV (c.1849A>G [p.Met617Val]) in the C2H2-type zinc finger (ZF) domain is a causative mutation for Skeletal defects, genital hypoplasia, and mental retardation (SGYMR)[6]. A SNV (c.1319G>A [p.Arg440Gln]) in the ZF domain was identified in brothers with ASD[7]. A SNV (c.1741A>T [p.Lys581*]) in the ZF domain was identified in a patient with schizophrenia[8, 9]. Other missense and nonsense mutations (highlighted in orange) in *ZBTB16* have been reported in ClinVar[10] (https://www.ncbi.nlm.nih.gov/clinvar), The Human Phenotype Ontology[11] (https://hpo.jax.org), and The Genome Aggregation Database[12] (gnomAD v2.1.1) (https://gnomad.broadinstitute.org) databases. From gnomAD v2.1.1, only the missense and nonsense mutations in the BTB/POZ and ZF domains are listed. Disorder-associated mutations are highlighted in magenta. Bold letters in black are mutations that may be associated with ASD. *BTB/POZ: BTB/ POZ domain for protein-protein interaction, C2H2-type zinc finger: Cys2-His2-type zinc finger domain for DNA binding, N: N-terminal, C: C-terminal, AA: amino acids.*

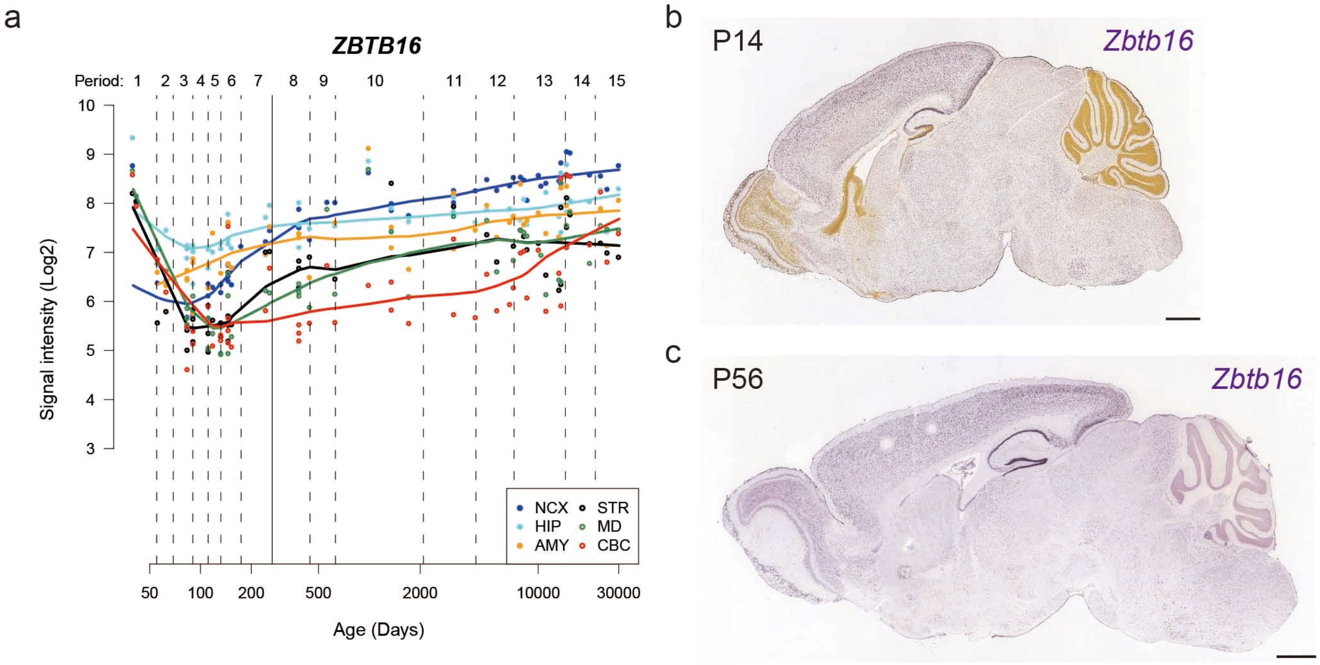


**Supplementary Figure 2. *Zbtb16* expression in the cortex.**
**a** *ZBTB16* expression trajectory in the neocortex, striatum, amygdala, hippocampus, midbrain, cerebellum from Human Brain Transcriptome (https://hbatlas.org; Kang et al. 2011). *NCX: neocortex, STR: striatum, AMG: amygdala, HIP: hippocampus, MD: midbrain, CBC: cerebellum.* **b, c** *Zbtb16* mRNA expression in sagittal sections of mouse brains at P14 (**b**) and P56 (**c**) detected by *in situ* hybridization. All images are from the Allen Brain Atlas. Scale bars: 1000 μm.

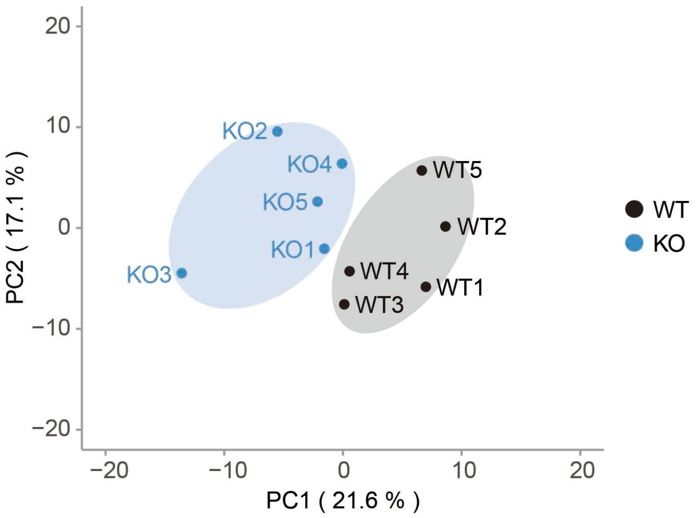


**Supplementary Figure 3. Principal component analysis (PCA) of *Zbtb16* genotypes.**
PCA in mouse prefrontal cortex, showing clear separation between WT and KO. *WT: wild-type, KO: Zbtb16 knockout.*

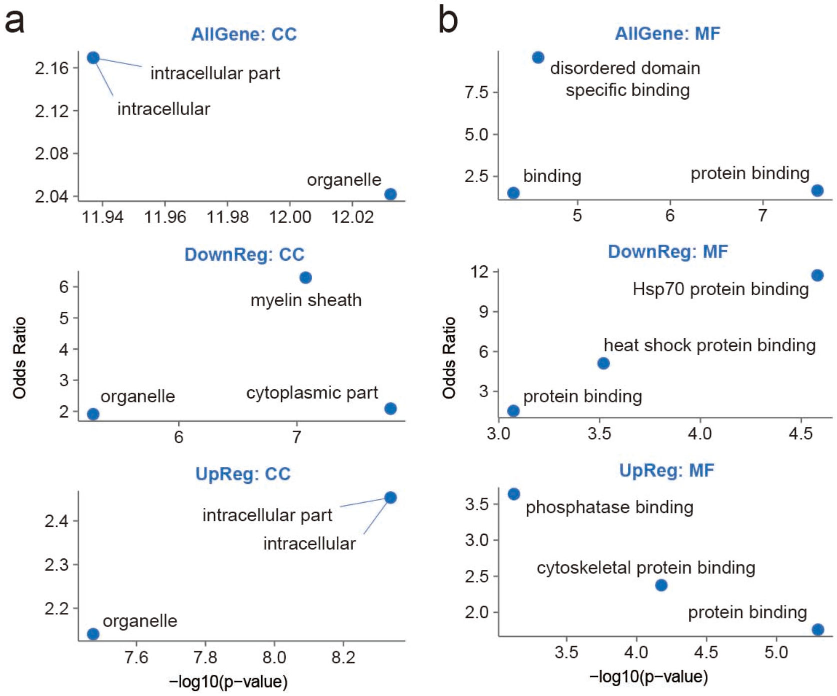


**Supplementary Figure 4. Gene ontology (GO) of *Zbtb16* differentially expressed genes (DEGs).** **a, b** GO analyses of *Zbtb16* DEGs in cellular component (CC) (**a**), and molecular function (MF) (**b**). Scatterplots represent the top 3 functions in each module. Y-axis=Odds Ratio, X-axis=-log10(p-value). *AllGene: all DEGs, DownReg: downregulated DEGs, UpReg: upregulated DEGs.*

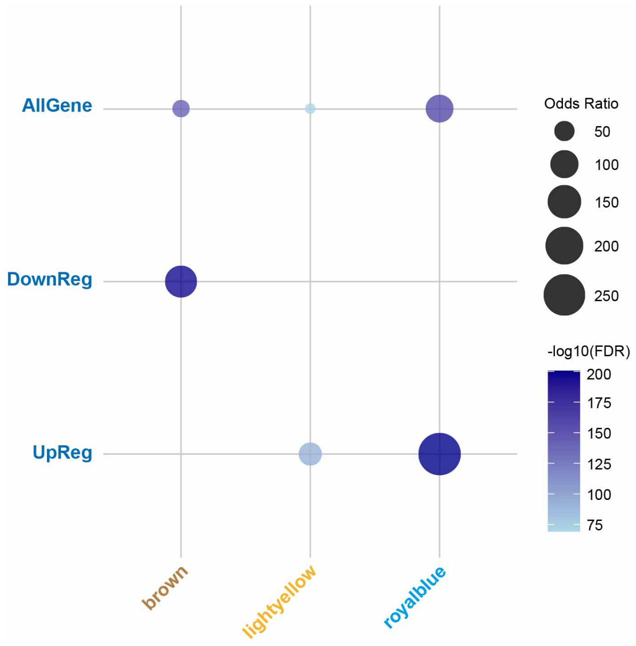


**Supplementary Figure 5. *Zbtb16* DEGs enrichments for *Zbtb16*-specific modules.** Downregulated *Zbtb16* DEGs are enriched in brown module. Upregulated *Zbtb16* DEGs are enriched in lightyellow and royalblue modules. *AllGene: all DEGs, DownReg: downregulated DEGs, UpReg: upregulated DEGs.*

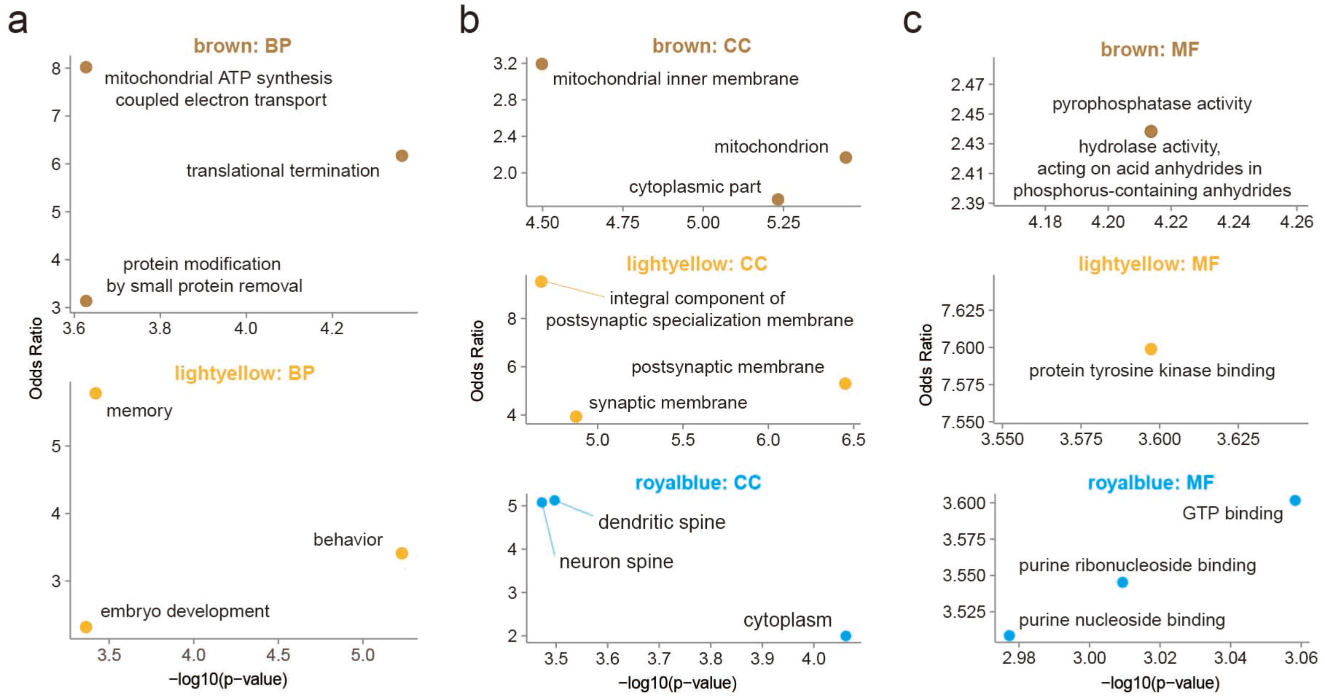


**Supplementary Figure 6. GO of *Zbtb16*-specific modules.**

**a-c** GO analyses of modules in biological process (BP) (**a**), CC (**b**), and MF (**c**). Scatterplots represent the top 3 functions in each module. Y-axis=Odds Ratio, X-axis=-log10(FDR).
